# Prediction of fast subcellular calcium signals based on label-free second harmonic generation microscopy measurements

**DOI:** 10.1101/2023.04.03.535276

**Authors:** Tihomir Georgiev

## Abstract

Calcium (Ca^2+^) sparks were predicted based on the label-free second harmonic generation (SHG) signal of myosin filaments with pix2pix, a conditional adversarial network. Predicted Ca^2+^ sparks and ground truth Ca^2+^ sparks show similar properties. Predicted Ca^2+^ spark frequencies and a predominant location at the periphery of skeletal muscle fibers were in good agreement with ground truth data. High frequencies of predicted Ca^2+^ sparks could be observed at y-shaped structures of myosin filaments. Important image pre-processing was applied in order to enhance the signal of Ca^2+^ sparks used for training. Only open-source software was applied. I provide the image pre-processing algorithm and several algorithms to evaluate the quality of predicted Ca^2+^ sparks. This technique can be extended to other label-free microscopy techniques and other fluorescent indicators developed during the last years. It remains a difficult task to predict the exact location of Ca^2+^ sparks.

## Introduction

Ca^2+^ is an important intracellular signal^1^. Ca^2+^ sparks are fast and small Ca^2+^ signals in different cell types^2,3^. I use the term Ca^2+^ sparks to describe localized Ca^2+^ signals in skeletal muscle fibers (Ca^2+^ sparks and Ca^2+^ bursts as defined in a previous publication^4^). High frequencies of Ca^2+^ sparks can be observed in permeabilized skeletal muscle fibers^5^ and during an osmotic treatment in intact skeletal muscle fibers^4^. Ca^2+^ sparks (short release events^6^) (38.9 ±1 ms)^4^ and Ca^2+^ bursts (events with extended openings^6^) (394.9±26.7 ms)^4^ were described in skeletal muscle fibers after hypotonic treatment^4^. I will specify if I use the term Ca^2+^ sparks to describe short localized Ca^2+^ signals. However, localized Ca^2+^ signals lasting more than a second have been described after osmotic treatment in skeletal muscle fibers^6,7^. Machine learning has been applied for the analysis of Ca^2+^ signals recently^8–14^. Generative adversarial networks have been introduced by Goodfellow et al^15^ and have been applied for segmentation of Ca^2+^ signals^16^. Machine learning approaches for the analysis of Ca^2+^ sparks are rare^17^. Recently, machine learning has been applied for segmentation and classification of Ca^2+^ sparks^17–19^. Additionally, T cell classification based on intracellular calcium signals has been carried out using machine learning predictions^20^.

In light microscopy, in silico labelling of subcellular structures based on bright field images has been achieved with deep learning^21,22^. Image translations of medical images have been generated with deep learning for different applications such as synthesis of post contrast MRI images^23^ or translation of MRI images to PET images^24^.

To my knowledge, this paper is the first attempt to translate structural images to images of Ca^2+^ signals. Second harmonic generation (SHG) microscopy is a nonlinear optical microscopy technique that allows the label-free visualization of myosin filaments in skeletal muscle fibers^25,26^. In vivo SHG measurements have been described in a previous study^27^. Localized Ca^2+^ signals have been investigated with respect to the myosin filaments in striated muscle cells previously ^28,29^. In this study, I used the SHG signal of myosin filaments as subcellular structural information to generate synthetic images of Ca^2+^ signals. The training of the network was carried out on pairs of images consisting of an SHG measurement and a two photon microscopy^30^ measurement of Ca^2+^ sparks in skeletal muscle fibers. For the training process of the network, an important image pre-processing of the Ca^2+^ signals was applied in order to enhance the signal of Ca^2+^ sparks compared to the cellular background. Different strategies have been applied to translate images previously^31,32^. I applied the pix2pix algorithm^32^ for the translation of images which is based on conditional adversarial networks. In principle this technique can be extended to other label-free microscopy techniques and other fluorescent indicators such as dopamine^33–36^, acetylcholine^37,38^, glutamate^39,40^, GABA^41^, ATP^14,42^, serotonin^43^, norepinephrine^44^ and adenosine^45^ indicators. The prediction of Ca^2+^ sparks based on subcellular structures is a difficult task since the exact location of Ca^2+^ sparks is difficult to predict. Additionally, Ca^2+^ sparks are small and fast subcellular signals in a noisy cellular environment.

Changes in the SHG signal depending on different conformations of myosin molecules have been described in skeletal muscle fibers and myofibrils^46–48^. An increase of the SHG signal in skeletal muscle fibers has been shown after an increase of the intracellular Ca^2+^ concentration^48^. Additionally, localized sarcomere contractions could be observed in response to Ca^2+^ sparks in cardiac muscle cells^29^. Therefore, I investigated possible subcellular changes of the SHG signal at regions of Ca^2+^ sparks.

## Results

### Generation of translated images

As a first step, Ca^2+^ sparks were enhanced compared to the background signal of the cell (Fig. 1a). Then, the network was trained on pairs of SHG images and enhanced Ca^2+^ images (Fig. 1b). The trained network was then applied to translate Ca^2+^ images based on SHG images (Fig. 1c). Fig. 1c shows only one Ca^2+^ spark in the translated image. Some translated Ca^2+^ sparks could be seen on every image of the measurement. Other translated Ca^2+^ sparks appeared only on some images of the measurement (Extended Data Fig. 1b, c, f). On several translated images two, three or four Ca^2+^ sparks could be observed (Extended Data Fig. 1a, b, c, f). Ca^2+^ sparks could be found at subcellular structures of the myosin pattern, so called y-shaped structures (Fig. 1d).

**Figure 1.**
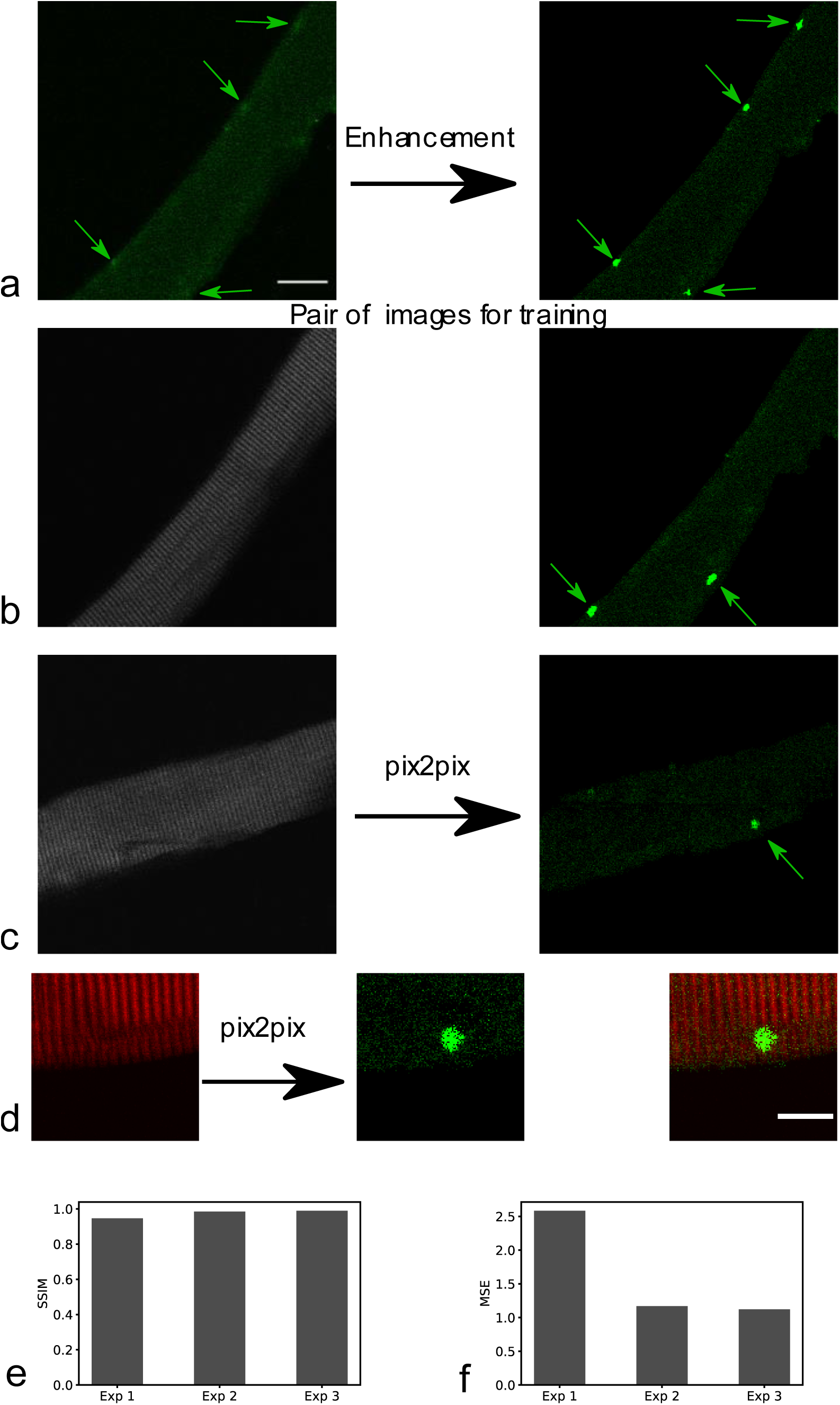
Generation of translated images. **a**, Pre-processing of Ca^2+^ sparks with enhancement of Ca^2+^ sparks compared to the background signal of the fiber. On the left side Ca^2+^ sparks (arrows) without enhancement are shown. On the right side Ca^2+^ sparks after enhancement can be seen. Scale bar 20 µm. **b**, Pairs of simultaneously measured SHG signal of the muscle fiber (left) and enhanced Ca^2+^ sparks (arrows right image) used to train the network. **c**, The trained network was applied to translate SHG images to Ca^2+^ images. The arrow indicates a Ca^2+^ spark in the translated image. Measurements of this fiber were not included in the training data set (Exp 1). **d**, Translation of SHG signal (red signal, left) to Ca^2+^ signal (green signal, centre) with the trained network. Overlay of both signals (right). Y-shaped structures can be seen in the SHG measurement (average of 5 images). Other measurements of this fiber and this plane were included in the training data set (Exp 3). Scale bar 10 µm. **e** and **f**, Structural similarity index (SSIM) and mean squared error (MSE) of the three experiments used to test the performance of the trained network.

I tested the performance of the trained network on three different experiments Exp 1 (experiment 1), Exp 2 (experiment 2), Exp 3 (experiment 3). In Exp 1 measurements of fibers were translated that were not included in the training data. Measurements showing a plane of a skeletal muscle fiber unseen during training were translated in Exp 2. However, measurements of another plane of this fiber (Exp 2) were included in the training data. Exp3 contains two measurements of a different fiber. Other measurements of the same plane of this fiber were included in the training data.

Structural similarity index (SSIM)^49^ and mean squared error (SME)^50^ were calculated (Fig. 1e and f). The translated images achieved high SSIM values.

### Properties of translated Ca^2+^ images

As a next step, I investigated several properties of translated images. The fiber masks of ground truth and predicted images were similar (Fig. 2a, b, c and d). The mean fiber intensity was lower in the predicted images, this was particularly the case in experiment 1 (Exp 1) (Fig. 2e). However, the mean of the fiber mean fluorescence intensity in the ground truth data of Exp 1 was higher than in the other two experiments and in the training data set.

**Figure 2.**
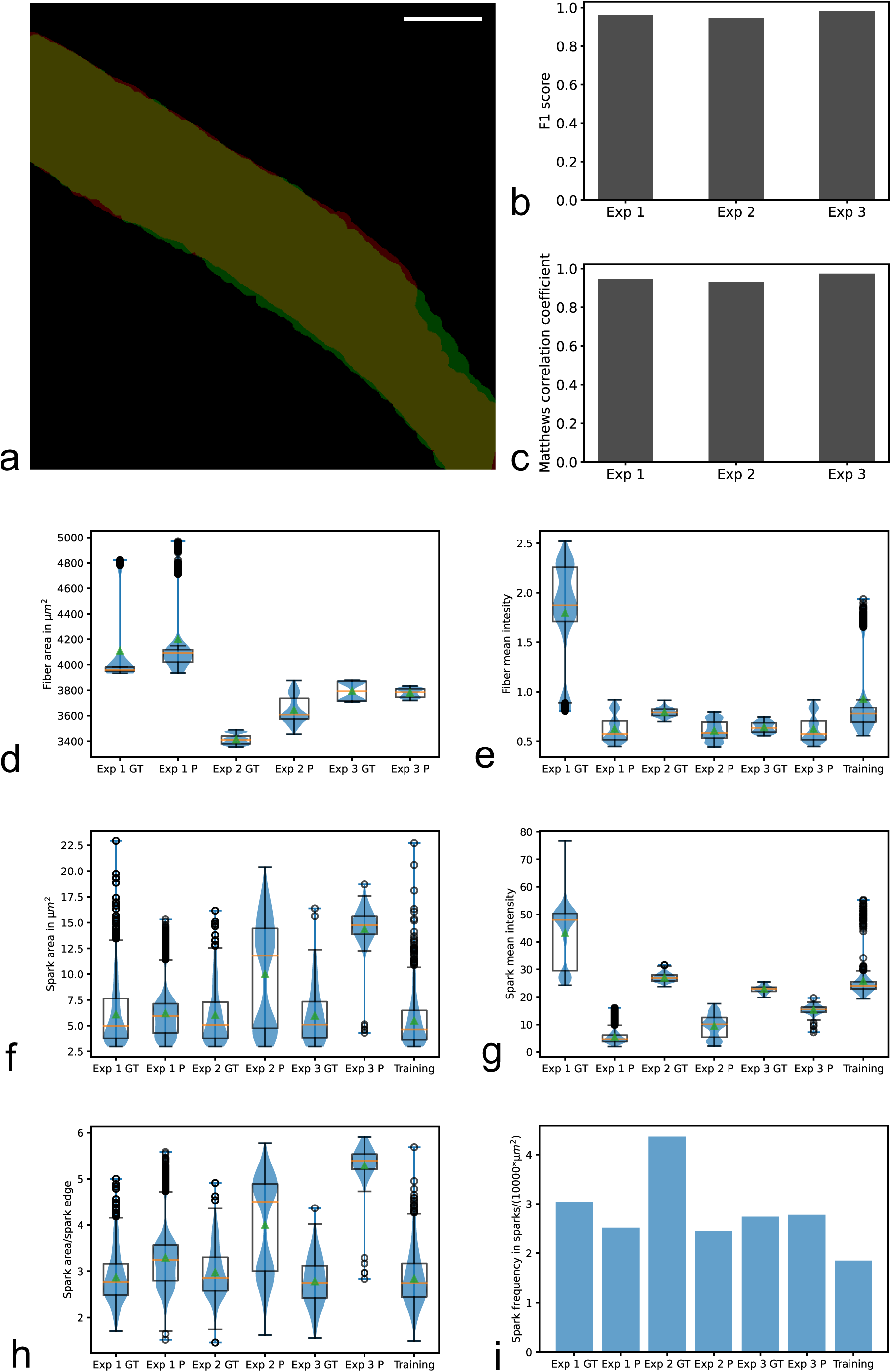
Properties of translated images of Ca^2+^ sparks. **a**, Ground truth (red) and predicted (green) fiber mask of a fiber unseen during training of the network (Exp 1). The overlay of both masks is yellow. Scale bar 20 µm. **b** and **c**, F1 scores and Matthews correlation coefficients for the evaluation of predicted fiber masks for the three experiments. **d**, **e**, **f**, **g** and **h**, Box plots and violin plots of the fiber area, fiber mean intensity, spark area, spark mean intensity and spark area/spark edge of the ground truth (Exp GT) and predicted (Exp P) images in the three experiments and the training data set. Orange lines are the medians of the data and green triangles are the means. **i**, Ca^2+^ spark frequencies of the ground truth and predicted images of the three experiments and the training data set.

Then, I investigated the properties of predicted Ca^2+^ sparks compared to ground truth Ca^2+^ sparks. I found that predicted Ca^2+^ spark area in Exp 1 was in good agreement with ground truth Ca^2+^ spark area (Fig. 2f). Interestingly, larger differences could be observed for the other two experiments. The predicted spark mean intensity was lower in all experiments (Fig. 2g). This was most pronounced in Exp 1. The predicted ratio Ca^2+^ spark area/ Ca^2+^ spark edge was larger than this ratio in the ground truth data for all experiments suggesting that predicted Ca^2+^ sparks are more circular (Fig. 2h).

The predicted Ca^2+^ spark frequency in Exp 1 of 2.52 sparks/(10000*µm^2^) was in good agreement with the ground truth Ca^2+^ spark frequency of 3.05 sparks/(10000*µm^2^) (Fig. 2i). The predicted Ca^2+^ spark frequency in experiment 3 (Exp 3) of 2.78 sparks/(10000*µm^2^) was even closer to the corresponding ground truth Ca^2+^ spark frequency of 2.74 sparks/(10000*µm^2^). The training data set had a Ca^2+^ spark frequency of 1.85 sparks/(10000*µm^2^). However, there was a larger difference in experiment 2 (Exp 2) between the predicted Ca^2+^ spark frequency of 2.46 sparks/(10000*µm^2^) and the ground truth Ca^2+^ spark frequency of 4.36 sparks/(10000*µm^2^).

### Translation of subcellular structures to fast subcellular signals

As a next step, I investigated specific subcellular regions of the muscle fiber known to have high frequencies of Ca^2+^ sparks, the fiber periphery^4^ and y-shaped structures of the myosin pattern^28^. Additionally, I investigated the ratio Ca^2+^ spark area over myosin filaments/spark area.

As expected, most of the predicted and ground truth Ca^2+^ sparks were located at the fiber periphery (Fig. 3a, b, and c). The ground truth Ca^2+^ spark frequency at the fiber periphery in Exp 1 showed a lower fraction of Ca^2+^ sparks compared to the predicted fraction. This difference can be also seen in the ratio Ca^2+^ spark area over fiber periphery/spark area (Extended Data Fig. 3b). One of the fibers used in Exp 1 showed a high frequency of Ca^2+^ sparks in the centre of the fiber not seen for the other five fibers (Extended Data Fig. 2). Interestingly, predicted images of this fiber showed several Ca^2+^ sparks in the fiber centre in close proximity to the ground truth Ca^2+^ sparks but not at such high frequencies (Extended Data Fig. 2). If this fiber was excluded from the analysis, there is a good agreement between ground truth data and predicted data (Extended Data Fig. 3a).

**Figure 3.**
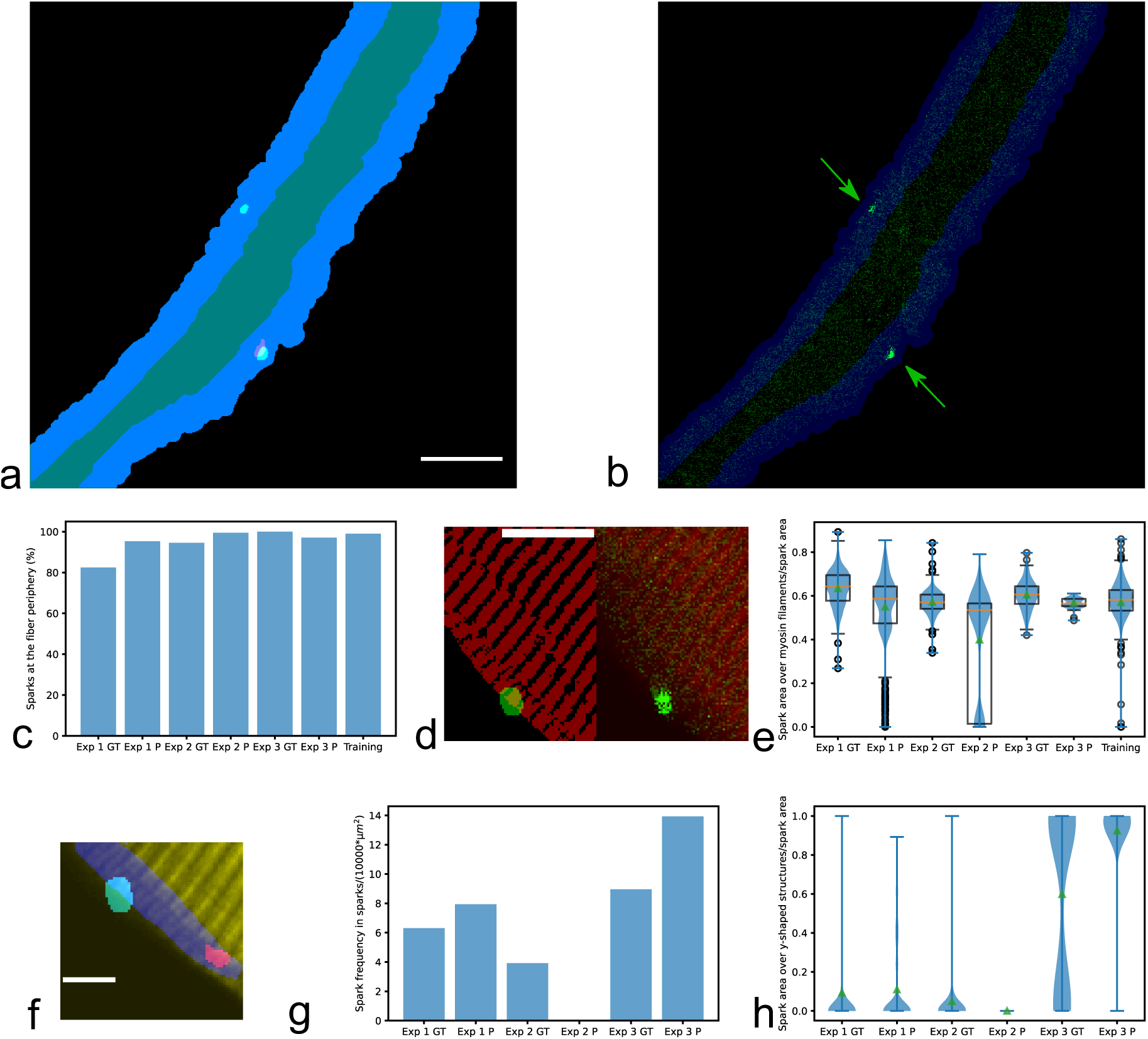
Translation of subcellular structures to fast subcellular signals. **a**, Binary images of the fiber centre (cyan), periphery of the fiber (blue) and Ca^2+^ sparks (green) based on a predicted Ca^2+^ image of a fiber unseen during training of the algorithm (Exp 1). Additionally, a binary image of a Ca^2+^ spark measured in the ground truth image is presented (red). Scale bar 20 µm. **b**, Overlay of the predicted Ca^2+^ signal (green) of the fiber shown in **a** and fiber periphery (blue). The arrows indicate Ca^2+^ sparks. **c**, Percentage of Ca^2+^ sparks at the fiber periphery for ground truth (Exp GT) and predicted (Exp P) images in the three experiments and the training data set. **d**, Overlay of binary images (left) of the myosin filaments (red) and a predicted Ca^2+^ spark. The binary SHG signal is based on a mean SHG signal of the whole measurement. Overlay of the mean SHG signal of the whole measurement (red) and the predicted Ca^2+^ signal (green) of the same image (right). This fiber was unseen during the training process of the algorithm (Exp 1). Scale bar 10 µm. **e**, Box plots and violin plots of the ratios of the Ca^2+^ spark area over myosin filaments and the Ca^2+^ spark area for ground truth (Exp GT) and predicted (Exp P) images in the three experiments and the training data set. Orange lines are the medians of the data and green triangles are the means. **f**, Overlay of the mean SHG signal during the whole measurement (yellow), mask of y-shaped structures (blue), binary image of a predicted Ca^2+^ spark (green) and binary image of a ground truth Ca^2+^ spark (red). The fiber was unseen during the training process (Exp 1). Scale bar 5 µm. **g**, Ca^2+^ spark frequencies at y-shaped structures for ground truth (Exp GT) and predicted (Exp P) images in the three experiments. **h**, Violin plots of Ca^2+^ spark area over y-shaped structures/Ca^2+^ spark area for ground truth (Exp GT) and predicted (Exp P) images in the three experiments. Green triangles are the means.

Then, I looked at the ratio Ca^2+^ spark area over myosin filaments/spark area (Fig. 3d and e) and found that this ratio was lower for predictions. Several predicted Ca^2+^ sparks were found at regions without myosin filaments (Exp 1 P and Exp 2 P). Such Ca^2+^ sparks at myosin free regions can be found in the training data set.

As a next step, I investigated the translation of y-shaped structures of the myosin filament pattern to Ca^2+^ sparks (Fig. 3f, g and h and Extended Data Fig. 4). In Exp 1, the predicted Ca^2+^ spark frequency at y-shaped structures of 7.94 sparks/(10000*µm^2^) was 1.26 times higher than the ground truth Ca^2+^ spark frequency at y-shaped structures of 6.31 sparks/(10000*µm^2^). In Exp 2, no Ca^2+^ sparks were predicted at y-shaped structures while ground truth Ca^2+^ frequency at y-shaped structures was 3.92 sparks/(10000*µm^2^). In Exp 3, the predicted Ca^2+^ spark frequency at y-shaped structures of 13.93 sparks/(10000*µm^2^) was 1.56 times higher than the ground truth Ca^2+^ spark frequency at y-shaped structures of 8.95 sparks/(10000*µm^2^).

### Accuracy of Ca^2+^ spark prediction

Predicted Ca^2+^ sparks seen in all image translations of a measurement were several times associated with high ground truth Ca^2+^ spark activity in close proximity (Fig. 4a, b, c and d, Extended Data Movie 1 and 2). The predicted and ground truth Ca^2+^ sparks partly overlapped in these cases. However, the exact overlap was not high. This is reflected in low Pearson correlation coefficients, Matthews correlation coefficients and F1 scores (Fig. 4 e, f, g). The predicted Ca^2+^ sparks seen in all image translations of a measurement were in close proximity or at y-shaped structures of the myosin filaments (Extended Data Fig. 5).

**Figure 4.**
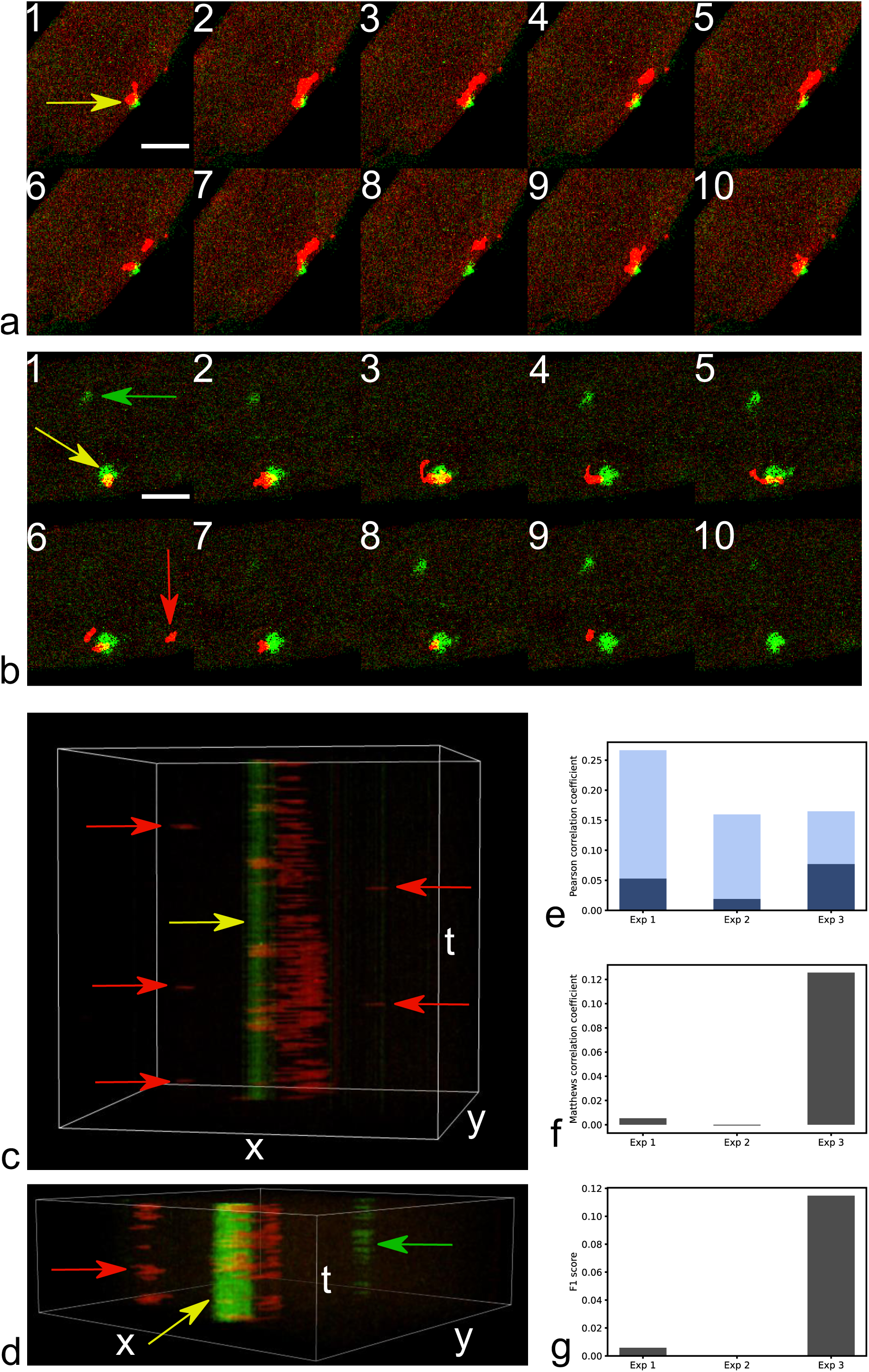
Accuracy of Ca^2+^ spark prediction. **a**, Image sequence (10 consecutive images) of the ground truth Ca^2+^ signal (red) and the predicted Ca^2+^ signal (green). The yellow arrow indicates a predicted Ca^2+^ spark. In close proximity to the predicted Ca^2+^ spark ground truth Ca^2+^ sparks can be observed. Predicted and ground truth Ca^2+^ sparks overlap partly sometimes. This fiber was unseen during the training process (Exp 1). Scale bar 10 µm. **b**, Image sequence of another fiber. Other measurements of this fiber and this plane were seen during the training process (Exp 3). The yellow arrow indicates a predicted Ca^2+^ spark (green) that is close to several ground truth Ca^2+^ sparks (red) and on several images they partly overlap. The green arrow shows at a predicted Ca^2+^ spark that is not in close proximity to ground truth signals. The red arrow indicates a ground truth Ca^2+^ spark that is not in close proximity to predicted Ca^2+^ sparks. Scale bar 10 µm. **c** and **d**, xyt volumes of the measurements partly shown as image sequences in **a** and **b**. Yellow arrows indicate predicted Ca^2+^ sparks in close proximity to many ground truth Ca^2+^ sparks partly overlapping with them. Red arrows show at ground truth Ca^2+^ sparks with no corresponding predicted Ca^2+^ sparks at this location. Green arrow shows at predicted Ca^2+^ sparks with no corresponding ground truth Ca^2+^ sparks at this location. Box volume 34.88 µm, 34.88 µm, 122.18 s in **c** and 34.88 µm, 34.88 µm, 40.18 s in **d**. **e**, Pearson correlation coefficients of the whole images (light blue) and of the overlapping fiber masks of predicted and ground truth images (dark blue) for the three experiments. **f** and **g**, Matthews correlation coefficients and F1 scores calculated based on binary images of ground truth Ca^2+^ sparks and predicted Ca^2+^ sparks for the three experiments.

Extended Data Fig. 6 and 7 present the results when only one fiber was included in Exp1.

### Does the algorithm translate subcellular structures or signals?

To investigate possible subcellular changes of the SHG signal at regions of Ca^2+^ sparks, I analysed if there is such correlation (Fig 5a, b and c). If there are Ca^2+^ dependent changes of the SHG signal, it is possible that the algorithm translates localized signals in SHG images to localized signals in Ca^2+^ images. The Pearson correlation coefficients for the different experiments are: Exp 1 GT 1nf: 0.0727, Exp 1 P 1nf: −0.4127, Exp 1 GT 1: −0.2998, Exp 1 P 1: −0.4564, Exp 1 GT 2: 0.0818, Exp 1 P 2: 0.6419, Exp 2 GT: 0.0299, Exp 2 P: −0.4498, Exp 3 GT: −0.1464, Exp 3 P: - 0.4887, Training: −0.1896.

**Figure 5.**
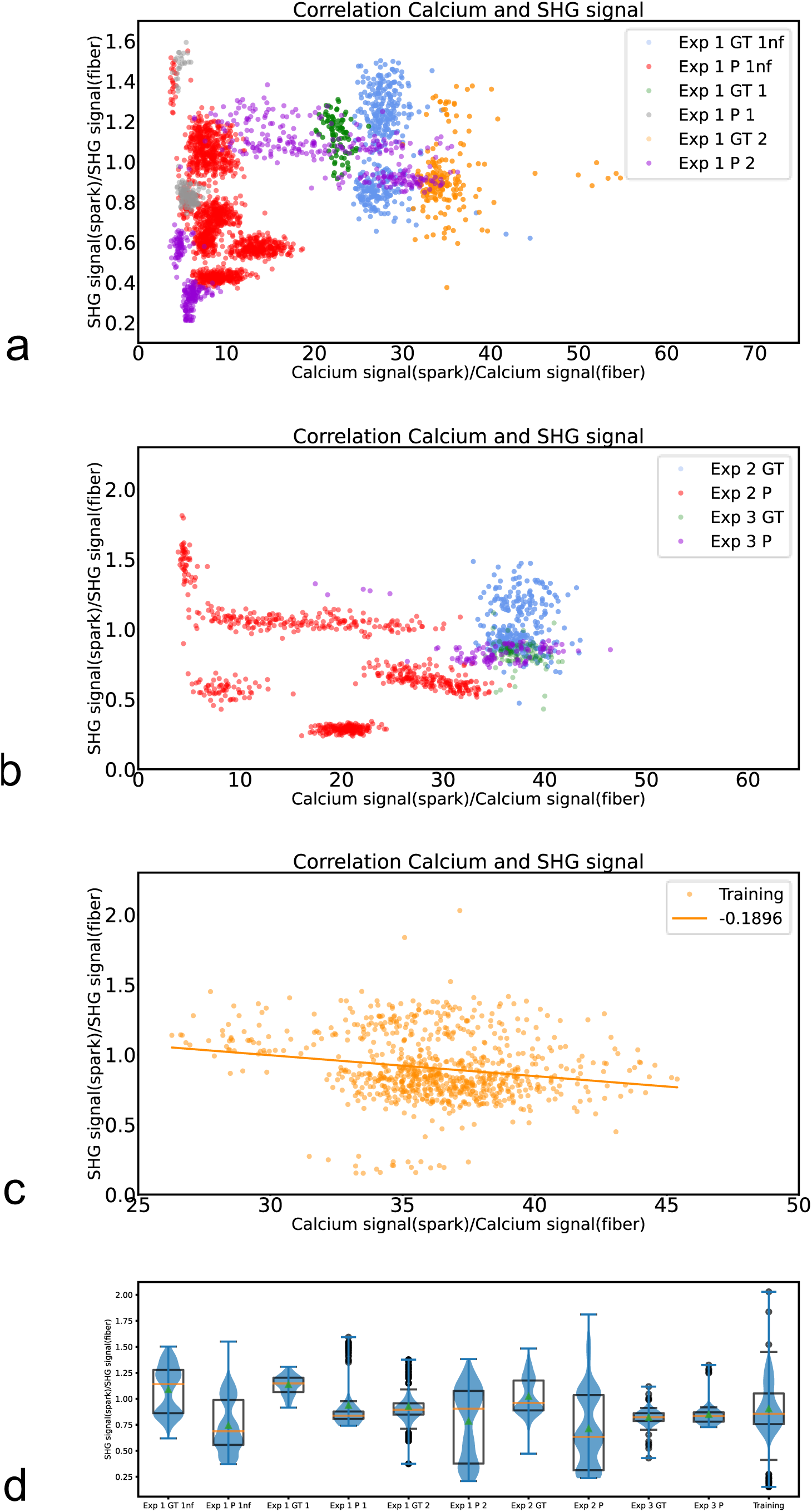
Does the algorithm translate subcellular signals to subcellular signals? Correlation of the mean SHG signal and Ca^2+^ signal at Ca^2+^ sparks divided by the mean SHG signal and the Ca^2+^ signal of the fiber, when regions of Ca^2+^ sparks were excluded from the fiber mask, for the three experiments (**a** and **b**) and the training data set (**c**). Exp1 (Experiment 1) fiber 1 was measured without a band pass filter in front of the SHG signal detector (Exp 1 GT 1nf and Exp 1 P nf (ground truth and predicted data)), another measurement of fiber 1 (Exp 1) was carried out with a band pass filter in front of SHG signal detector (Exp 1 GT 1 and Exp 1 P 1 (ground truth and predicted data)). Exp 1 GT 2 and Exp 1 P 2 are ground truth and predicted data of fiber 2 of Exp 1. Exp 2 GT and Exp 2 P and Exp 3 GT and Exp 3 P are ground truth and predicted data of Experiment 2 and 3. A linear regression is shown in **c** and the Pearson correlation coefficient is −0.1896 for the training data set. **d** Mean SHG signal at the region of a Ca^2+^ spark divided by the mean SHG signal of the fiber mask when regions of Ca^2+^ sparks were excluded from the fiber mask.

Interestingly, Exp 1 P 1nf shows negative Pearson correlation for the predicted data. The ratio SHG signal (spark)/SHG signal (fiber) is lower for Exp 1 P 1nf compared to Exp 1 GT 1nf (Fig 5d).

The F1 score for the fiber masks in Exp 1 c1nf (cell 1 without a band pass filter) is somewhat lower than the F1 score in Exp 1 c1 and Exp 1 c2 (Extended Data Fig.8). The F1 score for Ca^2+^ sparks is highest in Exp 1c1nf (Extended Data Fig.8).

I investigated if the SHG signal at Ca^2+^ sparks changes compared to the mean SHG signal of the measurement at the same Ca^2+^ sparks. 53.46% of Ca^2+^ sparks increased the SHG signal ratio>1 (1070 Ca^2+^ sparks, two mice, four fibers) (Fig. 6). The mean SHG signal ratio of Ca^2+^ sparks is not significantly changed 0.9996±0.0014 (mean±sem).

**Figure 6.**
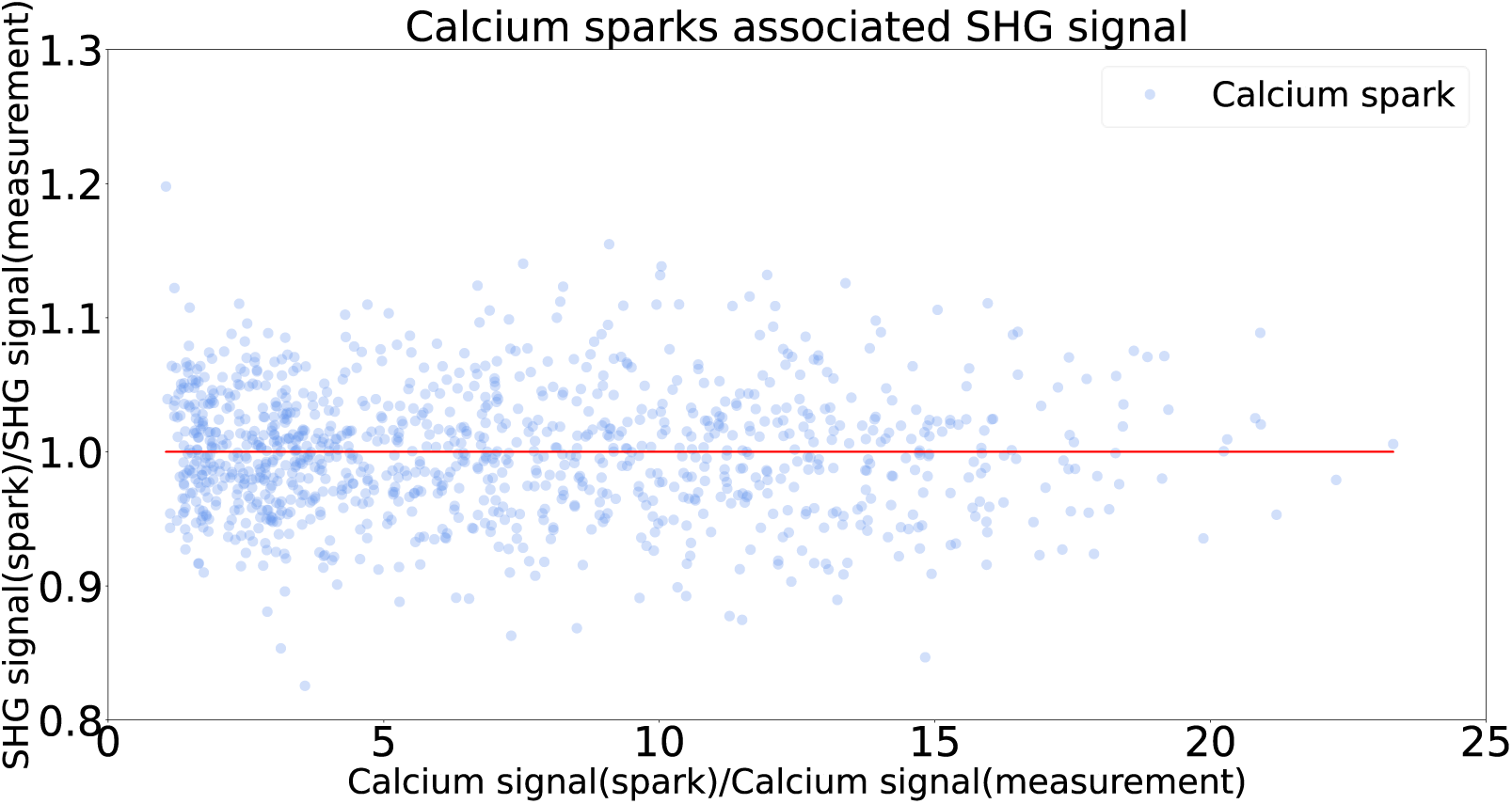
SHG signal changes associated with Ca^2+^ sparks. Ratios of SHG signal or Ca^2+^ signal at Ca^2+^ sparks and the mean SHG signal or the mean Ca^2+^ signal of the measurement (49 or 50 images) at the Ca^2+^ spark. Red line: Ratio SHG signal(spark)/SHG signal(measurement) of 1.

## Discussion

Several properties of Ca^2+^ signals predicted based on label-free SHG measurements were in good agreement with the ground truth Ca^2+^ signals. Predictions of fast subcellular signals based on measurements of label-free techniques have the potential to make obsolete elaborate staining of cells or one additional channel is then available for another fluorescent marker. The frequency of Ca^2+^ sparks, the fiber area and the Ca^2+^ spark area (in Exp 1) could be predicted quite well. Ca^2+^ sparks at myosin free regions might be nuclear Ca^2+^ sparks^7^. The poor prediction of Ca^2+^ sparks in Exp 2 suggests that a different plane of the same fiber is already a different scene with different myosin pattern and associated Ca^2+^ sparks signalling pattern. As expected, most predicted Ca^2+^ sparks were located at the fiber periphery. Small fiber shifts in part of the measurements might cause some inaccuracy in the analysis of Ca^2+^ sparks associated with y-shaped structures and the analysis of spark area over myosin filaments.

Predicted Ca^2+^ sparks seen in all images of a measurement were in several cases associated with locations of high ground truth Ca^2+^ sparks activity.

The better F1 scores for Ca^2+^ sparks in Exp 1 1nf compared to Exp 1 1 (Extended Data Fig. 8) might be explained by a different part of the fiber that is easier to predict. The influence of a different plane for the prediction of Ca^2+^ sparks has been shown in Exp 2. Although the training data shows a negative Pearson correlation, I think that results in Fig. 5 suggest that the algorithm translates subcellular structures. In contrast to the training data, Exp 1 GT 2 and Exp 2 GT show somewhat positive Pearson correlation. Additionally, a low SHG signal at Ca^2+^ sparks might be explained by specific positions within the fiber (perinuclear, nuclear signals or signals at the periphery of the fiber). Some Ca^2+^ sparks in skeletal muscle fibers might not be sufficient to increase the SHG signal as it was observed after a Ca^2+^ transient in skeletal muscle fibers^48^. Increased SHG signal has been observed for some sparks in the measurements presented here. But this needs further investigation and more sensitive optical systems might be able to measure Ca^2+^ sparks associated subcellular SHG signals more frequently. Measurements at most sensitive angles of relative laser polarization^47^ will facilitate the detection of such SHG signal changes. Better temporal resolution of the optical system will further improve the prediction of fast localized Ca^2+^ signals.

Such predictions might be more precise in cardiac muscle cells since local sarcomere contractions associated with Ca^2+^ sparks have been described in SHG measurements in a previous study^29^. Local changes in the sarcomere length as described in cardiac muscle cells^29^ may be an important information for the algorithm. However, I did not investigate such local changes in the sarcomere length. Additionally, polarization dependent effects of the SHG signal^46–48^ might influence the decision of the algorithm.

The prediction of the exact location of Ca^2+^ sparks based on subcellular structures remains a difficult task. In further improvement of the algorithm the 3D nature of xyt volumes should be taken into account instead of translating 2D images. Other existing algorithms such as U-net^51^ could be tested for such a task as it was done in an image translation study of medical images^23^. It can be expected that data augmentation of the training data set^14,51,52^, pre-processing of the SHG signal, denoising, larger data sets, powerful computational resources and algorithms designed for this specific task will further improve the accuracy of such image translations. In some cases and after further improvement of such predictions, label-free SHG measurements and a software similar to the one described in this manuscript might be used as a Ca^2+^ indicator in cardiac and skeletal muscle.

I hope that this work will stimulate further research to improve the translation of fast subcellular signals based on subcellular structures or other subcellular signals.

## Methods

### Data

The SHG images and two photon microscopy images used to train and test the algorithm were measured simultaneously in a previous publication^28^.

### Image pre-processing

The first step of the image pre-processing was to automatically identify Ca^2+^ sparks. Similar ideas for automatic detection of Ca^2+^ sparks have been described in a previous publication^53^. First, a gaussian filter was applied on each xyt measurement. Then, based on the gaussian signal image the mean intensity was calculated along the t axis, a fiber mask was generated using the mean image with a threshold of 2*σ, with σ the standard deviation. The mask was labelled and objects smaller than 3000 pixels were excluded. Then, a cleaned binary fiber mask was created. To exclude labelled structures outside the fiber, the Ca^2+^ measurement after application of a gaussian filter was multiplicated by the cleaned binary fiber mask. Binary image that includes the Ca^2+^ sparks was generated. Pixels were assigned a value of 1 if their value is higher than µ + 3.6* σ, with µ the mean and σ the standard deviation. µ and σ were measured within the fiber mask. Then objects of the binary images were labelled and objects smaller than 55 pixels were excluded. Then again, a binary image of Ca^2+^ sparks was created based on the cleaned labelled image.

Finally, the Ca^2+^ sparks in the original image were enhanced 10 times compared to the fiber background by reducing the fiber background 10 times. This step was important in order to make it easier for the network to see the Ca^2+^ sparks.

### Translation of Ca^2+^ sparks based on SHG images

19 measurements of 4 fibers from two mice were used to train the algorithm. The training was carried out with 4123 images containing 256×256 pixels with 818 Ca^2+^ sparks. Thereby, 1031 images containing 512×512 pixels were cropped to 4123 256×256 pixel images.

To test the performance of the trained network, three different experiments Exp 1 (experiment 1), Exp 2 (experiment 2), Exp 3 (experiment 3) were carried out.

Exp 1 consists of 545 512×512 pixel images from 5 measurements of two fibers from two mice (the same animals as used for training but different fibers), which were not included in the training data. 683 ground truth Ca^2+^ sparks are included in this data set.

Exp 2 consists of 196 512×512 pixel images from 4 measurements of a fiber that was included in the training data but showing a different plane. This data set contains 292 ground truth Ca^2+^ sparks.

Exp 3 consists of 98 512×512 pixel images from two measurements of a fiber. Different measurements of this muscle fiber were included at the same plane in the training data. The data set contains 102 ground truth Ca^2+^ sparks.

Fibers were selected if they showed clear Ca^2+^ sparks activity and a band pass filter was placed in the SHG signal detection path^28^ and no or little shift of the muscle fiber was observed during measurement. Additionally, 2 measurements were included in Exp 1 without a band pass filter in front of the SHG detector in order to test if a contamination of the SHG signal influences the prediction of Ca^2+^ sparks.

Since there is a high variability of Ca^2+^ sparks signalling patterns and because no augmentation was carried out during the training process, images in Exp 1 and Exp 2 were rotated at 90°, 180°, 270° and 360°. Therefore, the algorithm was tested on 2180 512×512 pixel images in Exp1 and 784 512×512 pixel images in Exp 2.

All images in the three experiments were cropped to 256×256 pixel images as input for the image translation algorithm.

In Exp 1 and in Exp 2 for the four angles 2307 and 702 Ca^2+^ sparks were predicted respectively. In Exp 3 103 Ca^2+^ sparks were predicted.

Additional analysis was carried out for Exp 1 (Fig. 5 and Extended Data Fig. 8). Exp 1 was divided in three different data sets. Exp 1 GT 1nf is the ground truth data set of fiber 1 without a band pass filter in front of the SHG signal detector. Exp 1 GT 1nf consists of 401 ground truth Ca^2+^ sparks. Exp 1 P 1nf are the predicted images of Exp 1 GT 1nf with 1628 Ca^2+^ sparks after rotation of images with 4 different angles as input. Exp 1 GT 1 is the ground truth data set of fiber 1 with a band pass filter in front of the SHG signal detector. This data set contains 90 Ca^2+^ sparks. The corresponding prediction Exp 1 P 1 for 4 rotations contains 155 Ca^2+^ sparks. Exp 1 GT 2 is the ground truth data set of fiber 2 from another mouse and contains 192 Ca^2+^ sparks. The corresponding prediction for 4 rotations contains 524 Ca^2+^ sparks.

The code for image translations was taken with minor modifications from https://www.tensorflow.org/tutorials/generative/pix2pix and is based on the publication of Isola et al^32^. The structure of the network was not modified. The network was trained for 35 epochs, batch size of 1.

### Analysis of translated images

As first analysis step of the translated images, fiber masks of the ground truth and predicted Ca^2+^ signal were generated in order to determine the fiber area and the mean fluorescence intensity within the fiber. First, a gaussian filter was applied on the enhanced Ca^2+^ images. Then, a binary image of the mask was created. Pixels were assigned the value 1 if their value was higher than 0.00001* σ, with σ the standard deviation of the corresponding image. In the enhanced ground truth Ca^2+^ image the Ca^2+^ signal outside the fiber was set to 0 as described previously in the image pre-processing part. In the predicted Ca^2+^ images some pixels with values higher than 0 could be rarely observed outside the fiber. To exclude these pixels, objects in the binary images satisfying the condition 0.00001* σ, were labelled. Then, objects smaller than 3000 pixels were excluded and a cleaned binary image was generated.

Predicted Ca^2+^ sparks were automatically identified as described in the image pre-processing section but with fiber masks generated as described in this section (analysis of translated images) since predicted images might show some variability of the generated fibers. Properties of Ca^2+^ sparks (Ca^2+^ spark area, Ca^2+^ spark mean fluorescence intensity and the ratio Ca^2+^ spark area/ Ca^2+^ sparks edges) were determined based on the labelled binary images of Ca^2+^ sparks. Ca^2+^ frequencies (densities) were defined as the following ratio: number of Ca^2+^ sparks in an experiment/ overall fiber mask area in an experiment.

An algorithm was developed to automatically detect the fiber periphery. First, the border of the image was detected using ndi.binary_delation of scipy. Then, pixels touching the image border and being part of the fiber mask were assigned a value of 1. Then, using a fiber erosion (ndi.binary_erosion of scipy) over 40 steps an image was iteratively created that shows the centre of the fiber. On each of the 40 steps the images with pixels assigned a value 1 when touching the image border and being part of the fiber mask were added to the eroded image in order to avoid the centre of the fiber touching the image border being assigned to the fiber border (512×512 pixel images Fig. 3a and b). With this eroded image and the fiber mask image an image of the fiber periphery was created. If there were overlapping pixels of the fiber periphery and the binary image of Ca^2+^ sparks, the Ca^2+^ spark was counted as a signal at the fiber periphery.

For the measurement of the Ca^2+^ spark area over myosin filaments, the mean SHG signal over the whole measurement was used. First, a gaussian filter was applied on the mean SHG signal. Then, a fiber mask was generated with pixels assigned the value 1 if they fulfil the condition > σ, with σ the standard deviation of the mean SHG signal. To create a binary image of the myosin filaments the module filters.rank.mean of scikit image was applied on the mean SHG signal with a disk size of 5. The images of the mean SHG signal were multiplied by the fiber mask. The Ca^2+^ spark area over the myosin filaments was calculated based on the binary image of the mean SHG signal and the labelled binary image of Ca^2+^ sparks.

Y-shaped structures were manually annotated on mean SHG measurements using Fiji^54^. With these binary images of y-shaped structures and the labelled binary images of Ca^2+^ sparks, the Ca^2+^ spark area over y-shaped structures was calculated. If 30% of the Ca^2+^ spark area overlaid y-shaped structures, it was counted as Ca^2+^ spark at y-shaped structures and the corresponding Ca^2+^ spark frequencies at y-shaped structures were calculated for the three experiments with the corresponding areas of y-shaped structures.

The ratios SHG signal(spark)/SHG signal(fiber) and Calcium signal(spark)/Calcium signal(fiber) in Fig. 5 are the mean SHG signal or Ca^2+^ signal at the region of a Ca^2+^ spark divided by the mean SHG signal or mean Ca^2+^ signal of the fiber mask when regions of Ca^2+^ sparks were excluded from the fiber mask.

The ratios SHG signal(spark)/SHG signal(measurement) and Calcium signal(spark)/Calcium signal(measurement) are the SHG signal and Ca^2+^ signal at the Ca^2+^ spark region divided by the mean SHG signal or mean Ca^2+^ signal over 49 or 50 images at the same region. Measurements were selected for analysis, if they showed no or negligible fiber shifts.

### Statistics

Pearson correlation coefficients and linear regression were calculated with Scipy^55^, Matthews correlation coefficients were calculated with Scikit learn^56^, structural similarity index (SSIM)^49^ and mean squared error^50^ (SME) were calculated with Scikit image^57^. Violin plots and box plots were plotted with Matplotlib^58^. Boxes in boxplots extend from Q1 to Q3 (with Q1 lower quartile and Q3 upper quartile), median is indicated by the horizontal line, triangles indicate means, upper and lower whiskers show last data point less than Q3 + 1.5*(Q3-Q1) and first data point greater than Q1-1.5 (Q3-Q1) with (Q3-Q1) the interquartile range and the circles indicate datapoints beyond the whiskers^58^. The F1 score was calculated as described by Moen et al^59^.

### Software

The following software was used in this paper: Python, Jupyter, Spyder, Tensorflow^60^, Tensorflow_io, Keras^61^, Scikit learn^56^, Scikit image^57^, Numpy^62^, Matplotlib^58^, Scipy^55^, Tifffile^63^, Os, Pathlib, Time, Datetime, IPython^64^, Xlsxwriter, ImageJ^65^, Fiji^54^, ChimeraX^66^, https://www.tensorflow.org/tutorials/generative/pix2pix and https://github.com/embl-bio-it/image-analysis-with-python/blob/master/session-3to5/image_analysis_tutorial_solutions.ipynb.

## Code availability

The code and the checkpoint used in this paper is provided at https://github.com/TihomirGeorgiev3/SHGtoSpark_part_1 and https://github.com/TihomirGeorgiev3/SHGtoSpark_part_2

## Data availability

Data will be provided upon request. Some images will be provided at https://github.com/TihomirGeorgiev3/SHGtoSpark_part_1 and https://github.com/TihomirGeorgiev3/SHGtoSpark_part_2. The algorithms can be tested on these images.

## Extended Data

**Extended Data Figure 1.**
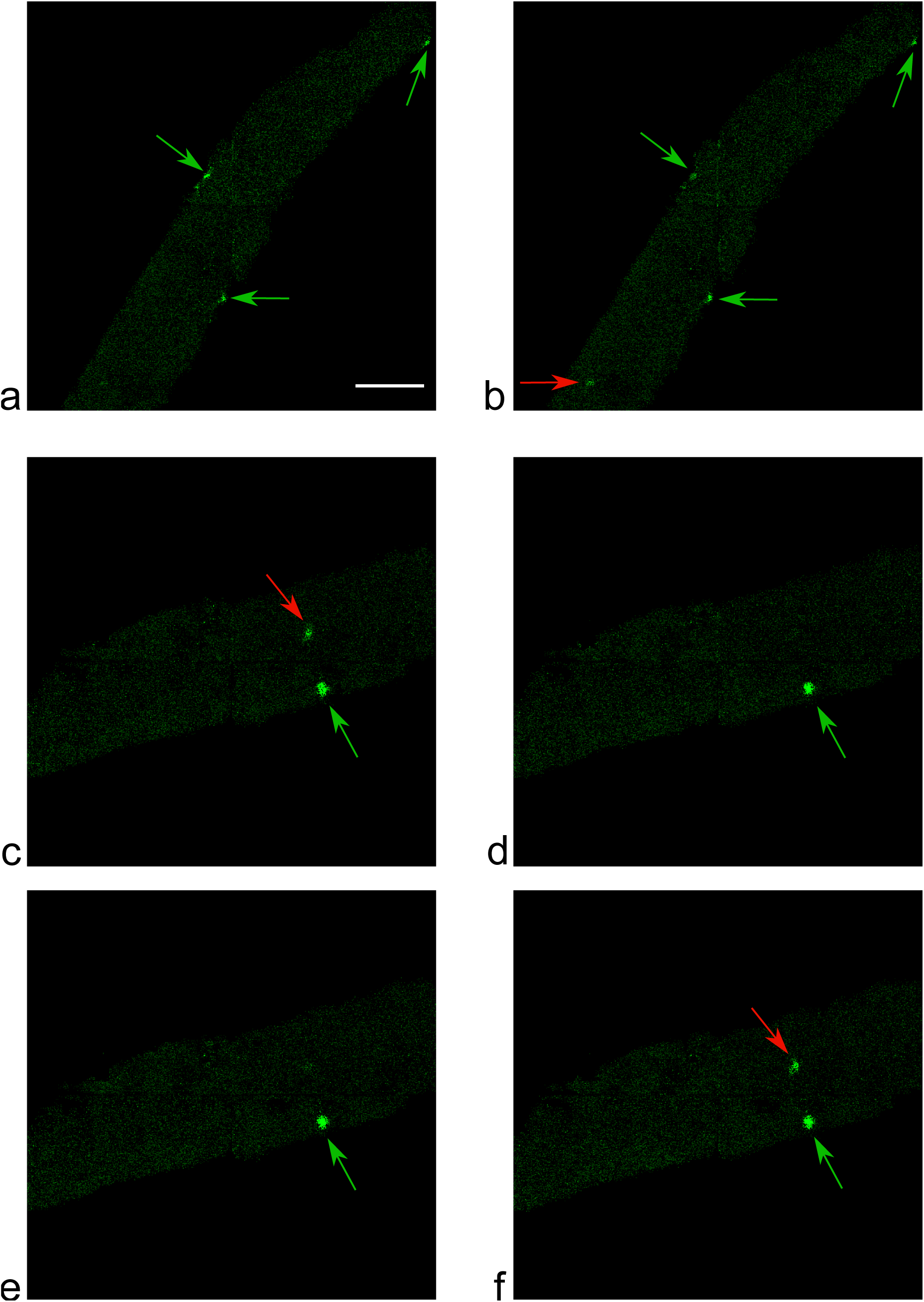
Predicted Ca^2+^ signalling patterns. **a** and **b**, Translations of two consecutive SHG images in a fiber not included in the training data set (Exp 1). Green arrows show at predicted Ca^2+^ sparks that can be seen on both images. The red arrow indicates a predicted Ca^2+^ spark that can be observed only on one of the images. Scale bar 20 µm. **c – f**, Translations of four consecutive SHG images of another fiber in Exp 1. Green arrows indicate predicted Ca^2+^ sparks seen in all images. Red arrows show predicted Ca^2+^ sparks seen only on two images and located more in the centre of the fiber.

**Extended Data Figure 2.**
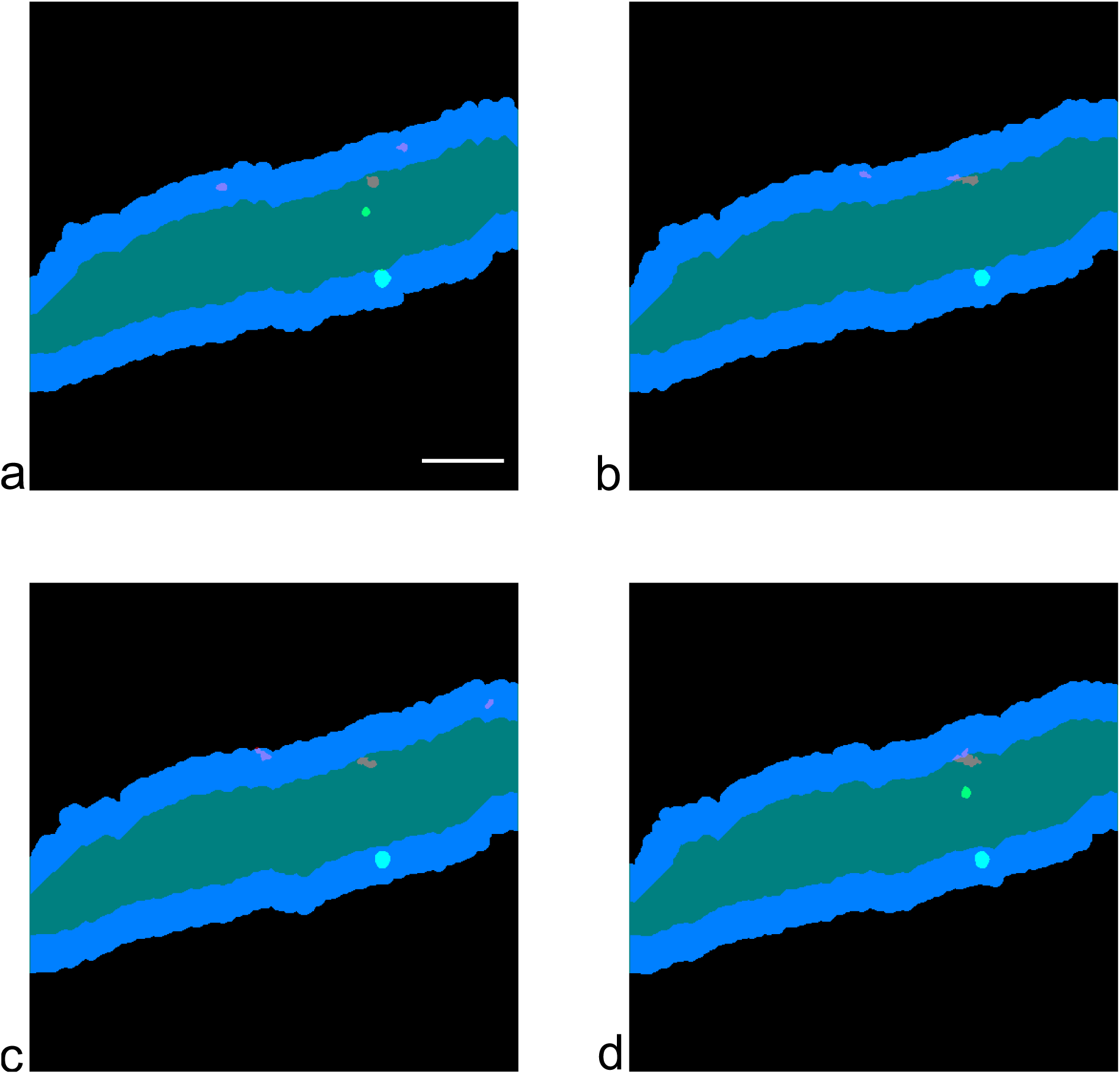
Predicted Ca^2+^ sparks at the centre of the fiber. Binary images of the predicted fiber periphery (blue), predicted fiber centre (cyan), predicted Ca^2+^ sparks (green) and ground truth Ca^2+^ sparks (red) of the four images shown in Extended Data Fig. 1c – f. In **a** and **d**, two predicted Ca^2+^ sparks at the centre of the fiber can be observed. In **a** and **c**, two ground truth Ca^2+^ sparks at the centre of the fiber can be seen. The predicted and ground truth Ca^2+^ sparks are located in close proximity. In **c**, a small part of one of the three ground truth Ca^2+^ sparks extends beyond the predicted fiber mask. This was seen extremely rarely since there is a high agreement between predicted and ground truth fiber masks. Scale bar 20 µm.

**Extended Data Figure 3.**
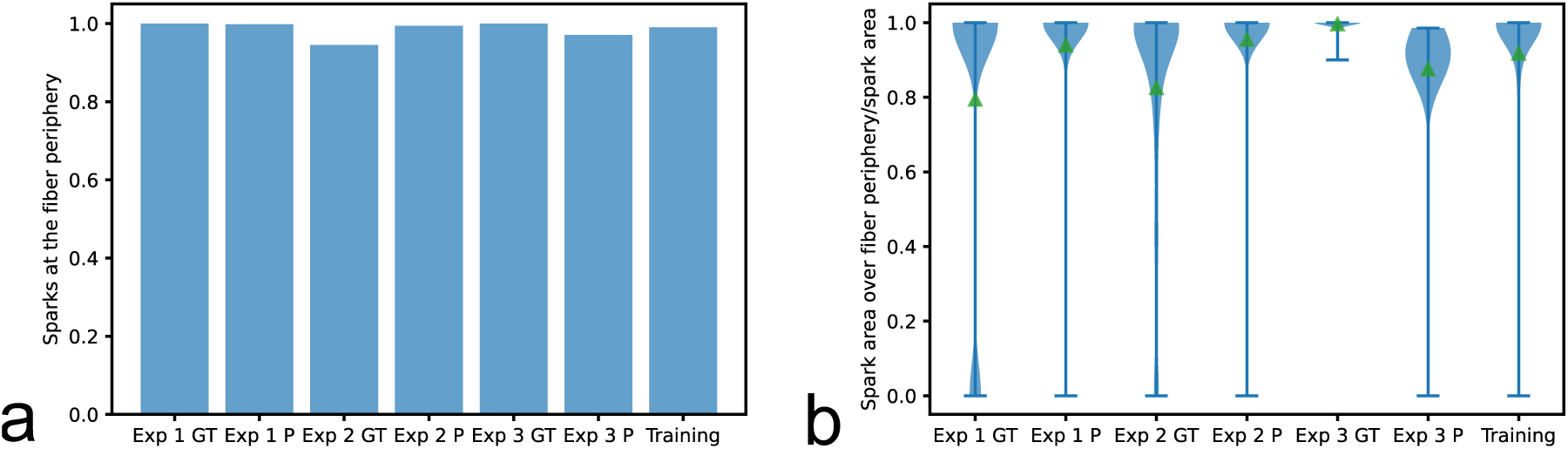
Ca^2+^ sparks at the fiber periphery. **a**, Fraction of Ca^2+^ sparks at the fiber periphery when one of two fibers in Exp1 was excluded from the analysis. The excluded fiber showed an unusually high Ca^2+^ spark frequency at the center of the fiber. **b**, Violin plots of the ratio of Ca^2+^ spark area over the fiber periphery and Ca^2+^ spark area for the three experiments (GT means ground truth and P prediction) and the training data set. Green triangles are the means.

**Extended Data Figure 4.**
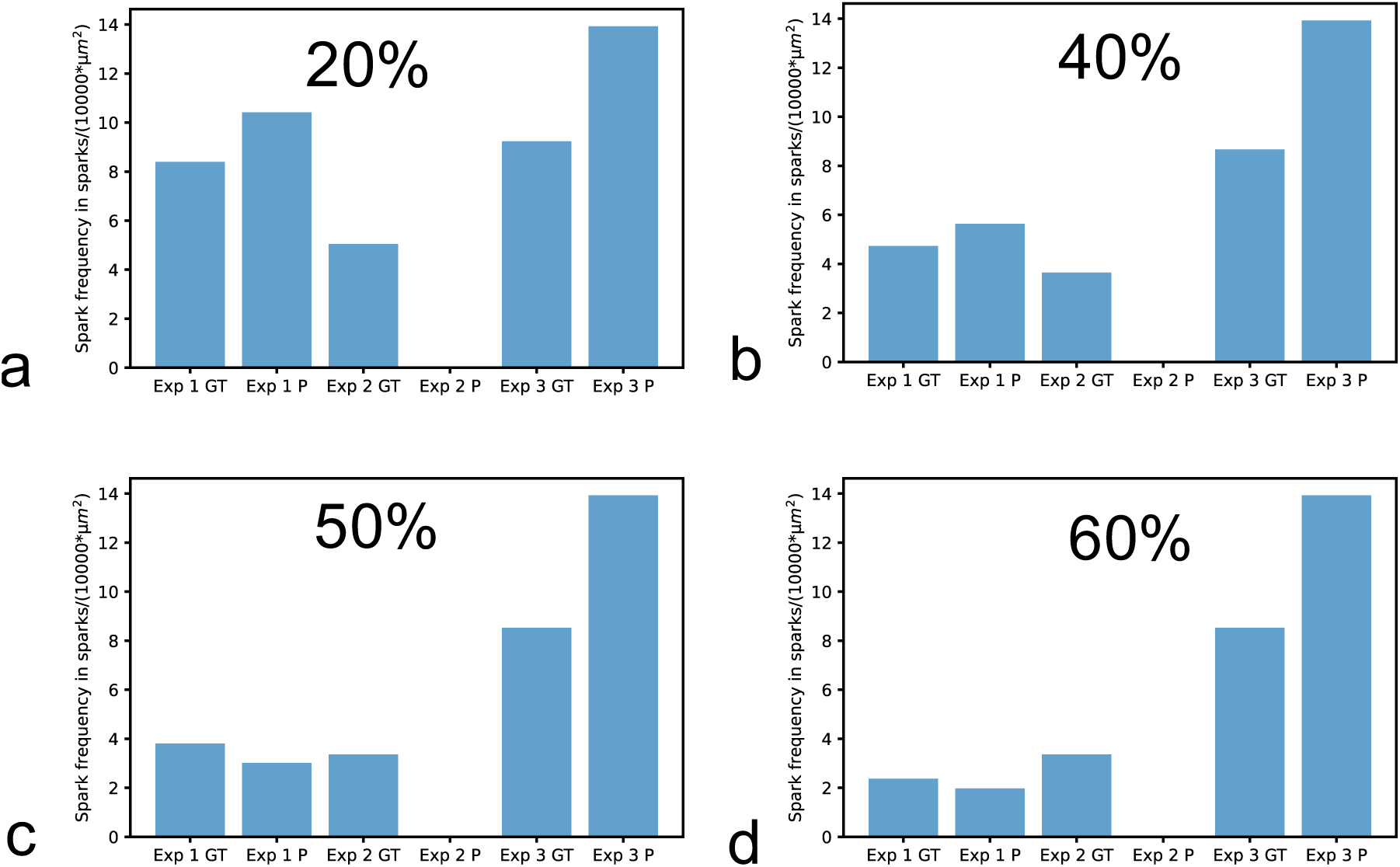
Ca^2+^ spark frequencies at y-shaped structures. **a – d**, Ca^2+^ spark frequencies for the three experiments. Ca^2+^ sparks were counted as Ca^2+^ sparks at y-shaped structures if at least 20% (**a**), 40% (**b**), 50% (**c**) and 60% (**d**) of the Ca^2+^ spark area covered y-shaped structures.

**Extended Data Figure 5.**
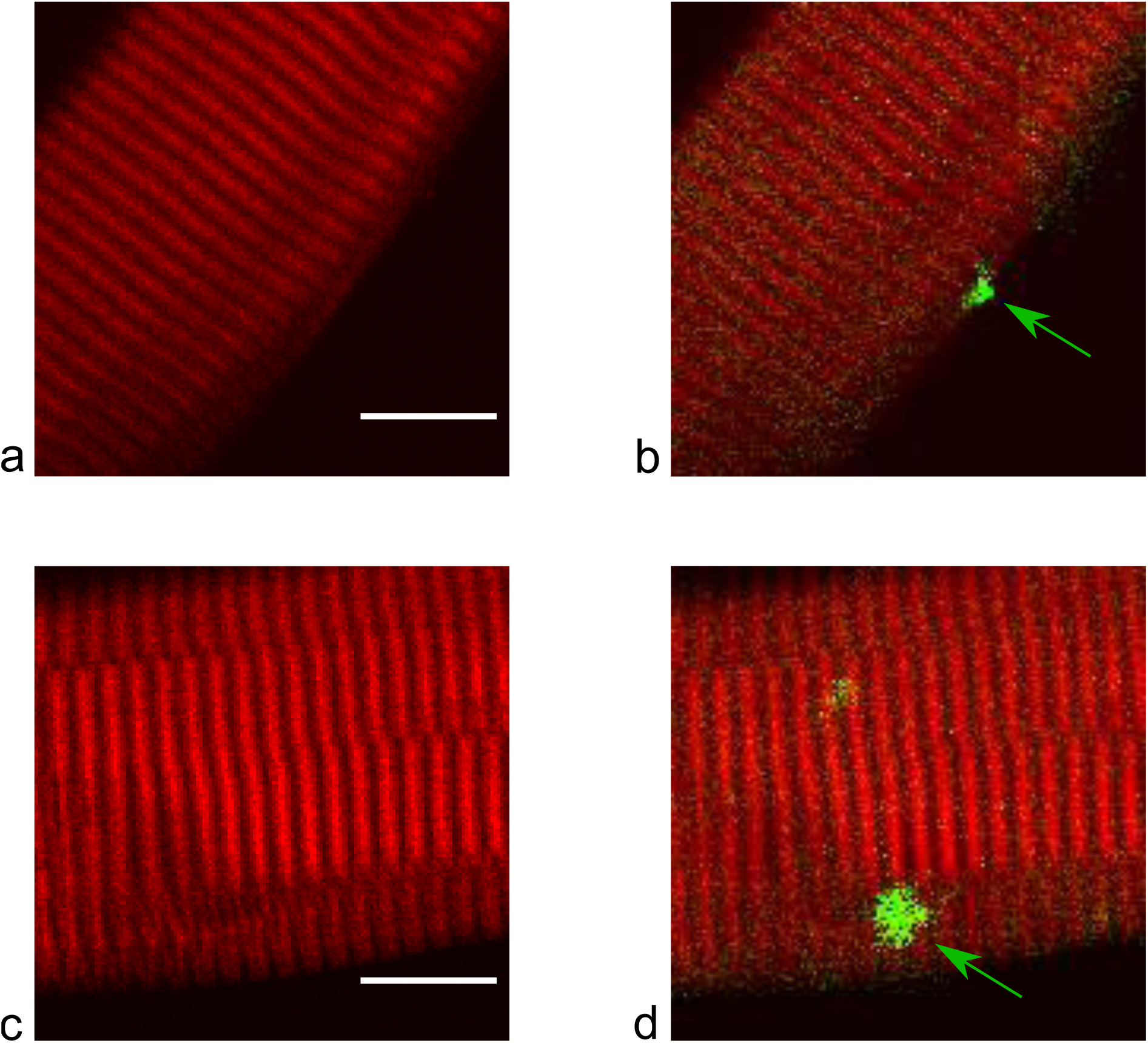
Ca^2+^ sparks seen in all image translations of a measurement associated with y-shaped structures. **a**, SHG measurement of the fiber shown in Fig. 4a. Scale bar 10 µm. **b**, Overlay of the predicted Ca^2+^ signal (green) (first image in Fig. 4a) and the SHG signal (red). The arrow indicates a predicted Ca^2+^ spark that could be seen in all image translations of this measurement (149 images) and is located in close proximity to y-shaped structures. **c**, SHG measurement of the fiber shown in Fig. 4b. Scale bar 10 µm. **d**, Overlay of the predicted Ca^2+^ signal (green) (first image in Fig. 4b) and the SHG signal (red). The arrow shows a predicted Ca^2+^ spark that could be seen in all image translations of this measurement (49 images) and is located at y-shaped structures.

**Extended Data Figure 6.**
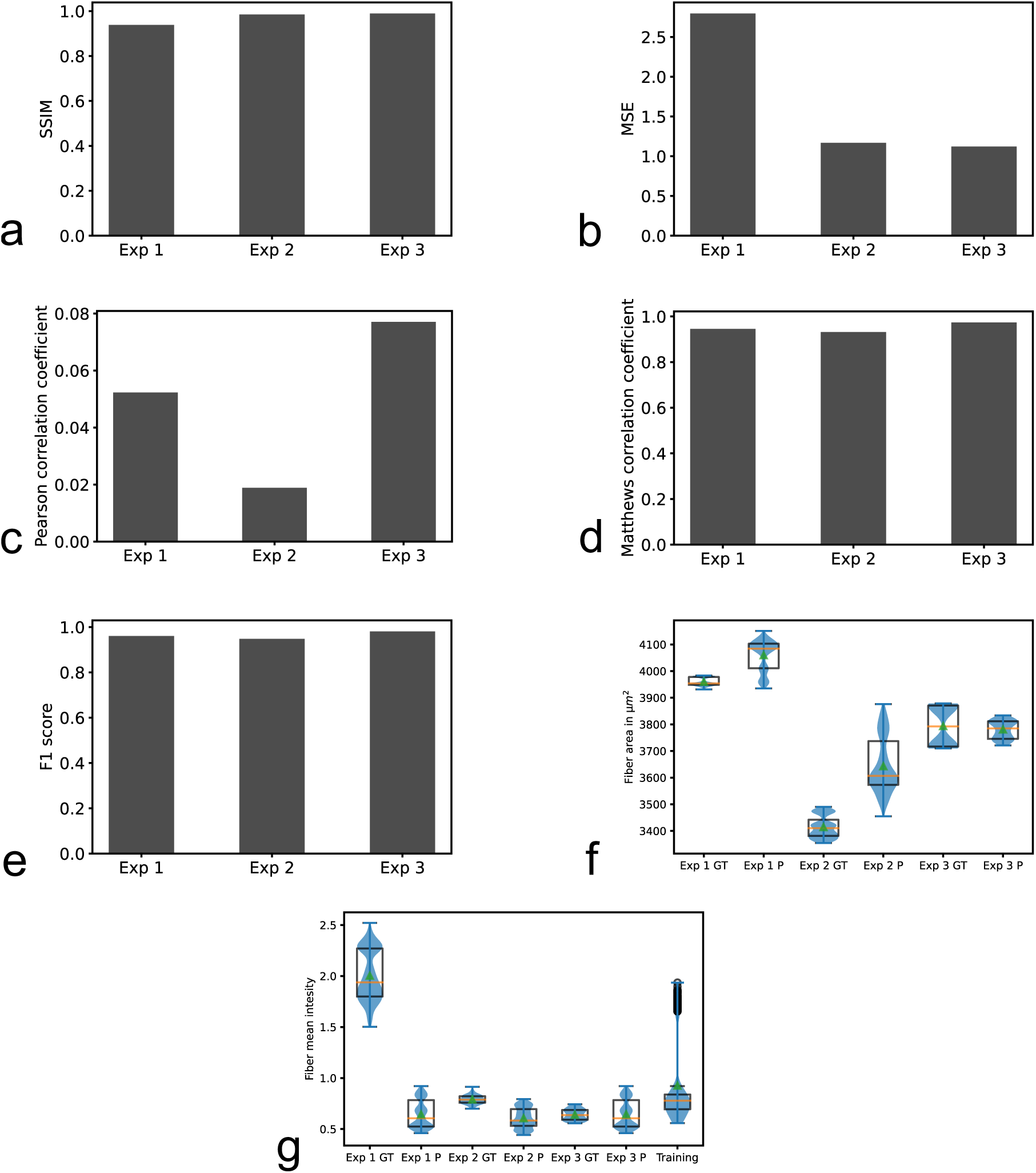
Properties of generated images with only one fiber in Exp1. Only one fiber in Exp1 was included in the analyses presented in this figure. This fiber was shown in Fig. 2a, in Fig. 3a, b, d, f and in Fig. 4a, c. 1788 images out of 2180 images in Exp 1 are measurements of this fiber. **a** and **b**, Structural similarity index (SSIM) and mean squared error (MSE) of the three experiments. **c**, Pearson correlation coefficients of the overlapping fiber masks of predicted and ground truth images for the three experiments. **d** and **e**, Matthews correlation coefficients and F1 scores calculated based on ground truth and predicted fiber masks for the three experiments. **f**, Box plots and violin plots of fiber areas of ground truth (GT) and predicted (P) fiber masks for the three experiments. **g**, Box plots and violin plots of fiber mean intensity of ground truth (GT) and predicted (P) fiber masks for the three experiments and the training data set.

**Extended Data Figure 7.**
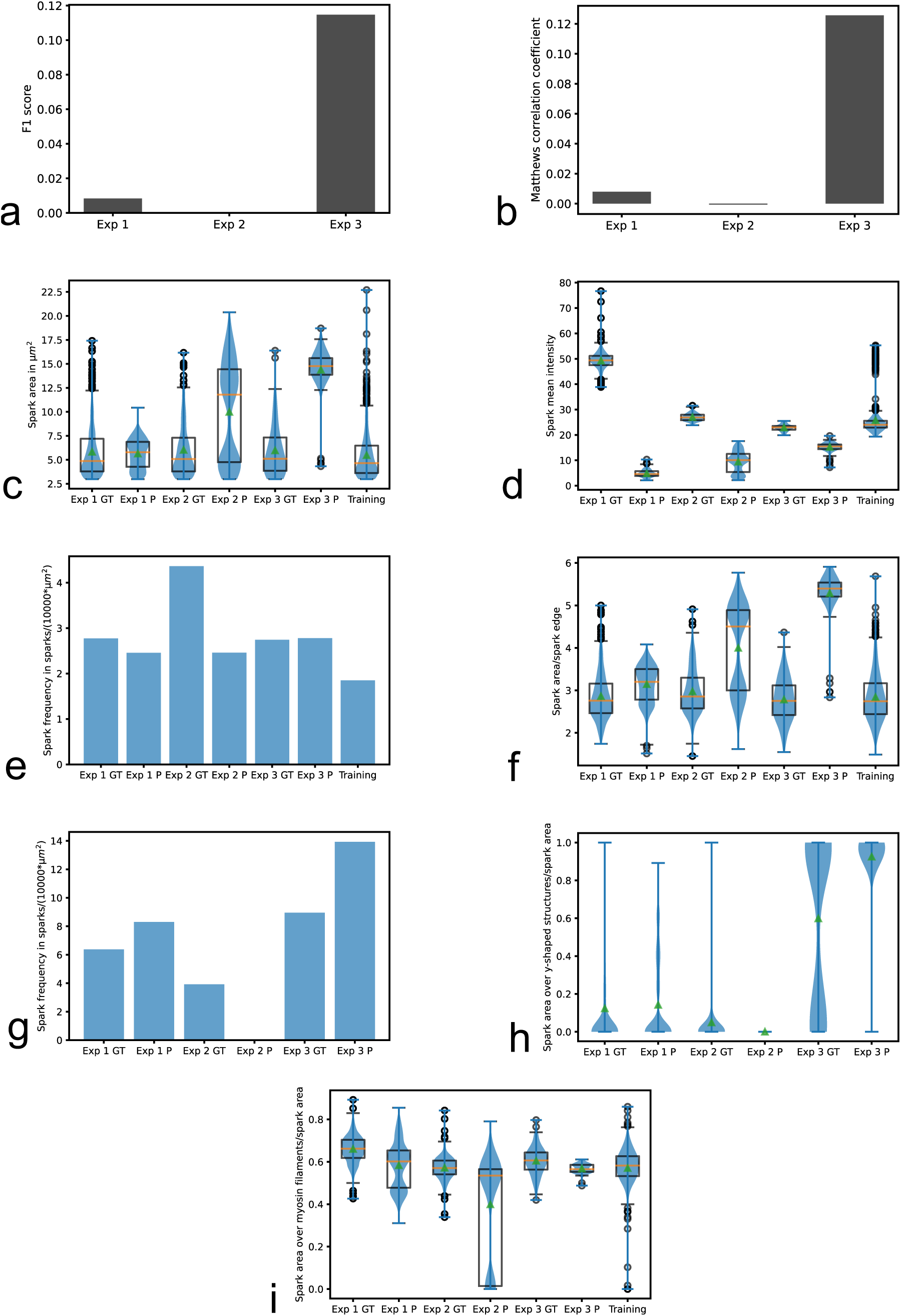
Properties of predicted Ca^2+^ sparks with only one fiber in Exp1. Only one fiber in Exp1 was included in the analyses presented in this figure. This fiber was shown in Fig. 2a, in Fig. 3a, b, d, f and in Fig. 4a, c. 1788 images out of 2180 images in Exp 1 are measurements of this fiber. **a** and **b**, F1 scores and Matthews correlation coefficients for the three experiments calculated based on binary images of ground truth and predicted Ca^2+^ sparks. **c** and **d**, Box plots and violin plots of Ca^2+^ spark area and Ca^2+^ spark mean intensity for the three experiments (GT means ground truth and P prediction) and the training data set. **e**, Ca^2+^ spark frequencies of the three experiments and the training data set. The ground truth Ca^2+^ spark frequency of Exp1 was 2.77 sparks/(10000*µm^2^). The predicted Ca^2+^ spark frequency of Exp1 was 2.46 sparks/(10000*µm^2^). **f**, Box plots and violin plots of the ratios Ca^2+^ spark area/Ca^2+^ spark edge for the three experiments and the training data set. **g**, Ca^2+^ spark frequencies at y-shaped structures for the three experiments. The ground truth Ca^2+^ spark frequency at y-shaped structures of Exp1 was 6.37 sparks/(10000*µm^2^). The predicted Ca^2+^ spark frequency at y-shaped structures of Exp1 was 8.30 sparks/(10000*µm^2^). **h**, Violin plots of Ca^2+^ spark area over y-shaped structures/Ca^2+^ spark area for the three experiments. **i**, Box plots and violin plots of the ratios Ca^2+^ spark area over myosin filaments/Ca^2+^ spark area for the three experiments and the training data set.

**Extended Data Figure 8.**
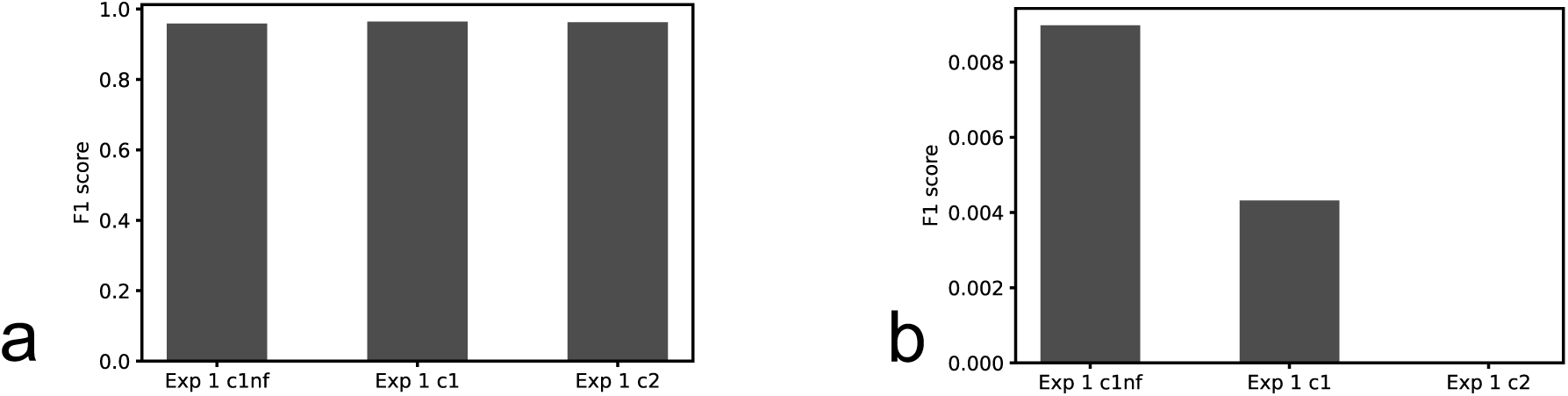
F1 scores for binary images of the Ca^2+^ fiber mask (a) and binary images of Ca^2+^ sparks (b) in Exp 1. Fiber 1 (mouse 1) measured without a band pass filter in front of the SHG signal detector (Exp 1 c1nf), fiber 1 (mouse 1) measured with a band pass filter in front of the SHG signal detector (Exp 1 c1) and fiber 2 (mouse 2) with a band pass filter in front of the SHG signal detector (Exp 1 c2).

**Movie 1**

xyt volume shown in Fig 4c. Box volume 34.88 µm, 34.88 µm, 122.18s.

**Movie 2**

xyt volume shown in Fig 4d. Box volume 34.88 µm, 34.88 µm, 40.18s.

